# Large scale genome-wide association study reveals that drought induced lodging in grain sorghum is associated with plant height and traits linked to carbon remobilisation

**DOI:** 10.1101/865667

**Authors:** Xuemin Wang, Emma Mace, Yongfu Tao, Alan Cruickshank, Colleen Hunt, Graeme Hammer, David Jordan

## Abstract

Sorghum is generally grown in water limited conditions and often lodges under post-anthesis drought, which reduces yield and quality. Due to its complexity, our understanding on the genetic control of lodging is very limited. We dissected the genetic architecture of lodging in grain sorghum through genome-wide association study (GWAS) on 2308 unique hybrids grown in 17 Australian sorghum trials over 3 years. The GWAS detected 213 QTL, the majority of which showed a significant association with leaf senescence and plant height (72% and 71% respectively). Only 16 lodging QTL were not associated with either leaf senescence or plant height. The high incidence of multi-trait association for the lodging QTL indicates that lodging in grain sorghum is mainly associated with plant height and traits linked to carbohydrate remobilisation. This result supported the selection for stay-green (delayed leaf senescence) to reduce lodging susceptibility, rather than selection for short stature and lodging resistance *per se*, which likely reduces yield. Additionally, our data suggested a protective effect of stay-green on weakening the association between lodging susceptibility and plant height. Our study also showed that lodging resistance might be improved by selection for stem composition but was unlikely to be improved by selection for classical resistance to stalk rots.

**Key message:** We detected 213 lodging QTL and demonstrated that drought induced stem lodging in grain sorghum is substantially associated with stay-green and plant height, suggesting a critical role of carbon remobilisation.

## Introduction

Water availability is a primary constraint to crop production worldwide. Among many cereal crops, sorghum is renowned for its tolerance to drought and heat (Hadebe et al. 2017). However, sorghum is often exposed to various types of water stress with drought during grain filling being common (Chapman et al. 2000; Kholová et al. 2013). One of the impacts of water stress during grain filling in sorghum is lodging. Lodging is the permanent displacement of crop stems from their vertical position. It can be caused either by buckling at a basal internode (stem lodging) or by root rotation in the soil (root lodging) (Pinthus 1974). Stem lodging is the major type of lodging induced by drought. When water stress occurs during grain filling, sorghum plants senesce and subsequently lodge due to stem collapse or breakage at the basal internodes (Henzell et al. 1984). This form of lodging is often observed in grain sorghum hybrids grown in Australia (Henzell et al. 1984), the USA (Rosenow 1977), and Argentina (Frezzi and Teyssandier 1980).

The occurrence and severity of lodging depends on the growth environment and the growth stage of the crop. Yield losses due to lodging in grain sorghum in production conditions are not well documented; however, it is believed that lodging, which occurs following water stress during the grain filling stage, causes the most grain loss world-wide, especially in regions where grains are harvested mechanically (Johnson et al. 1997).

The causes of lodging are not well understood. However, many factors may be associated with lodging, including stay-green, plant height, root and stalk rots, and stem characteristics.

Stay-green is an important trait providing tolerance to drought in many cereal crops including sorghum, wheat, maize, and rice (Cha et al. 2002; Christopher et al. 2008; Borrell et al. 2014a, b; Zhang et al. 2019). The stay-green phenotype is expressed as the delayed leaf senescence during grain filling. It is affected by many factors that influence the source-sink relationships within the plant (Borrell and Hammer 2000; Borrell et al. 2001), such as restricted tiller number and reduced upper leaf size resulting in reduced canopy size and pre-anthesis water demand (Borrell et al. 2014b), reduced grain number (Borrell and Hammer 2000), and possibly increased water uptake through narrow nodal root angle (Mace et al. 2012). Stay-green can increase water availability during post-anthesis water stress (Borrell et al. 2014a), which reduces the level of carbohydrate remobilisation by ensuring that photosynthesis continues, and consequently reduces the intensity of carbon remobilisation from the stems and leaves (Borrell and Douglas 1996).

Unlike stay-green, plant height not only affects carbohydrate supply/demand but also can have direct physical effects on lodging. Tall sorghum plants produce a higher stem biomass per grain than short crops (George-Jaeggli et al. 2011). This indicates that tall sorghum plants have more stem reserves available for translocation to the grains during post-anthesis water stress than short sorghum plants, which could potentially lead to less lodging in tall plants than in short plants. On the other hand, tall sorghum plants generally have higher yield than short sorghum plants (Jordan et al. 2003), and higher yielding sorghums are more susceptible to terminal drought stress than are lower yielding sorghums (Rosenow et al. 1983). Additionally the higher centre of gravity and greater leverage force in tall sorghums means that tall, high yielding genotypes are more prone to lodging under drought conditions after flowering.

This already complex interaction of factors is further complicated by the presence of soil borne root and stalk rotting pathogens, such as *Macrophomina phaseolina* and *Fusarium* spp., which may accelerate lodging. These fungi are necrotrophic in nature; while they may invade healthy plants, they are typically unable to colonise healthy tissue (Dodd 1980). However, when the stem begins to senesce under water stress during grain filling, they attack the dying roots and stalks (Srinivasa Reddy et al. 2008). The fungi secrete cell-wall-degrading enzymes (CWDE) to upregulate CWDE-encoding genes, degrading the integrity of the pith and rind in the basal internodes, which weakens the stalk and further exacerbates lodging (Bandara et al. 2018).

Lodging is also affected by stem mechanical properties. Associations between stem lodging and stem strength, rigidity, and chemical composition including lignin density have been observed (Bashford et al. 1976; Esechie et al. 1977; Murray et al. 2008; Gomez et al. 2017, 2018). Lignin is one of the key biomolecules in vascular plants and its functions include structural support and water transport (Boerjan et al. 2003; Higuchi 2006). Reduced lignin content has been shown to lead to increased lodging susceptibility in maize (Sun et al. 2018).

Because of its complex nature, limited progress has been made in understanding the genetic architecture of lodging in sorghum. To date, three studies (Kebede et al. 2001; Murray et al. 2008; Srinivasa Reddy et al. 2008) detected seven QTL associated with lodging in sorghum with no common QTL being identified across these studies. Given the complex set of traits known to affect lodging, the lack of QTL overlap is unsurprising and it is likely that the number of QTL affecting this trait is large.

The current study aims to dissect the genetic architecture of lodging and the pleiotropic effect of lodging QTL on stay-green and plant height in large-scale multi-environment field trials in Australia’s northern grain belt. We further investigate the co-localisation of lodging QTL with other traits including QTL for stalk rot disease resistance and genes in the lignin biosynthesis pathway.

## Materials and Methods

### Breeding Trials

This study made use of data from 17 trials of the Australia sorghum pre-breeding program run by the University of Queensland, the Department of Agriculture and Fisheries in Queensland, and the Grains Research and Development Corporation (UQ/DAF/GRDC). The trials were grown in a three year period from 2015-2017 and sampled the major sorghum growing region of Australia from northern New South Wales to central Queensland (from 22.295°S to 31.268°S; from 147.517°E to 152.105°E; Table 1). The 17 trials were selected from a larger set of 38 trials grown by the UQ/DAF/GRDC pre-breeding program during this period on the basis that lodging was recorded in the trial. Lodging is a critical trait for sorghum in Australia and is almost always scored whenever it occurs.

**Table 1.**
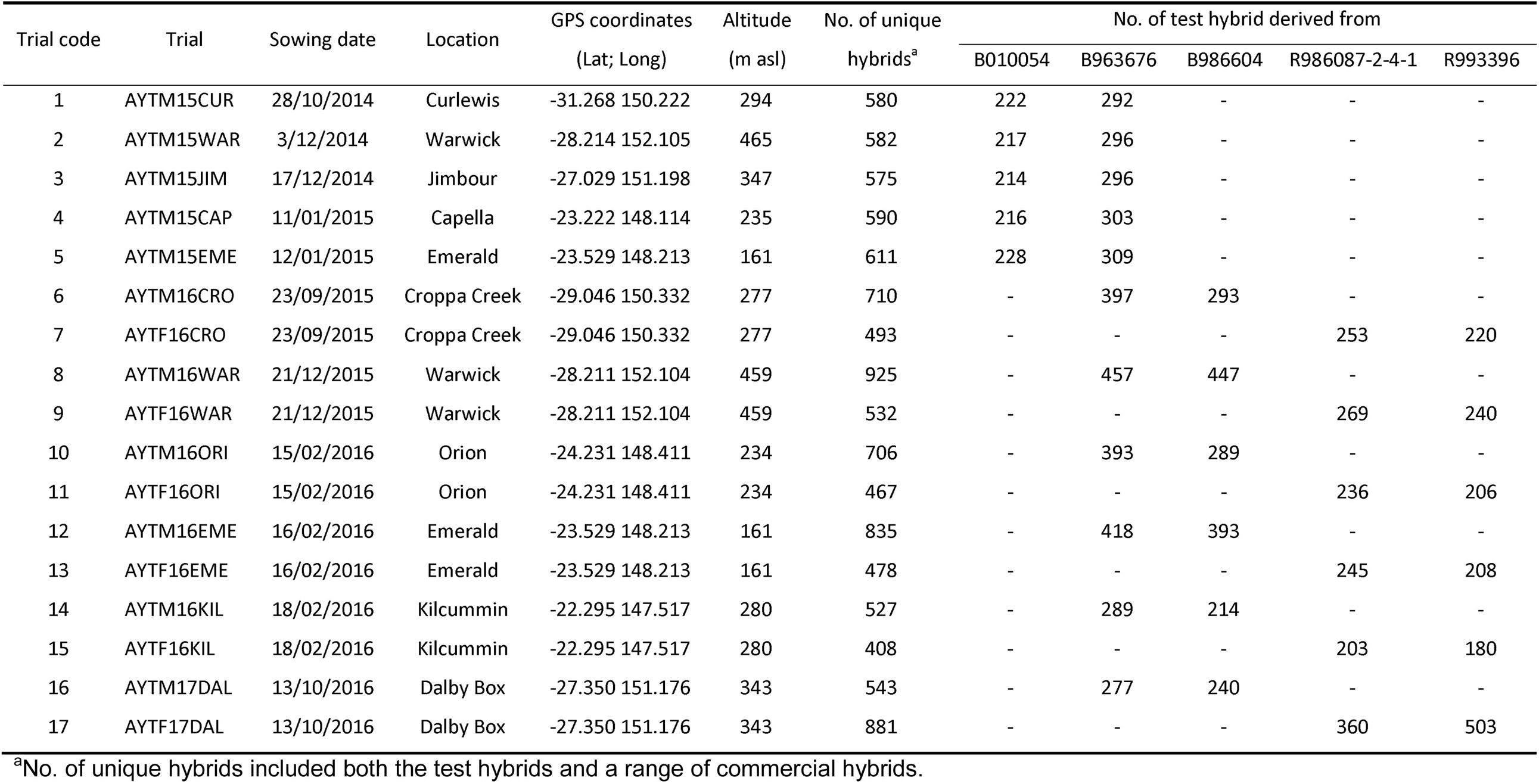
Description of the 17 field trials in the 2015-17 growing seasons.

| Trial code | Trial | Sowing date | Location | GPS coordinates<br>(Lat; Long) | Altitude<br>(m asl) | No. of unique<br>hybrids<sup>a</sup> | No. of test hybrid derived from |  |  |  |  |
| --- | --- | --- | --- | --- | --- | --- | --- | --- | --- | --- | --- |
|  |  |  |  |  |  |  | B010054 | B963676 | B986604 | R986087-2-4-1 | R993396 |
| 1 | AYTM15CUR | 28/10/2014 | Curlewis | -31.268 150.222 | 294 | 580 | 222 | 292 | - | - | - |
| 2 | AYTM15WAR | 3/12/2014 | Warwick | -28.214 152.105 | 465 | 582 | 217 | 296 | - | - | - |
| 3 | AYTM15JIM | 17/12/2014 | Jimbour | -27.029 151.198 | 347 | 575 | 214 | 296 | - | - | - |
| 4 | AYTM15CAP | 11/01/2015 | Capella | -23.222 148.114 | 235 | 590 | 216 | 303 | - | - | - |
| 5 | AYTM15EME | 12/01/2015 | Emerald | -23.529 148.213 | 161 | 611 | 228 | 309 | - | - | - |
| 6 | AYTM16CRO | 23/09/2015 | Croppa Creek | -29.046 150.332 | 277 | 710 | - | 397 | 293 | - | - |
| 7 | AYTF16CRO | 23/09/2015 | Croppa Creek | -29.046 150.332 | 277 | 493 | - | - | - | 253 | 220 |
| 8 | AYTM16WAR | 21/12/2015 | Warwick | -28.211 152.104 | 459 | 925 | - | 457 | 447 | - | - |
| 9 | AYTF16WAR | 21/12/2015 | Warwick | -28.211 152.104 | 459 | 532 | - | - | - | 269 | 240 |
| 10 | AYTM16ORI | 15/02/2016 | Orion | -24.231 148.411 | 234 | 706 | - | 393 | 289 | - | - |
| 11 | AYTF16ORI | 15/02/2016 | Orion | -24.231 148.411 | 234 | 467 | - | - | - | 236 | 206 |
| 12 | AYTM16EME | 16/02/2016 | Emerald | -23.529 148.213 | 161 | 835 | - | 418 | 393 | - | - |
| 13 | AYTF16EME | 16/02/2016 | Emerald | -23.529 148.213 | 161 | 478 | - | - | - | 245 | 208 |
| 14 | AYTM16KIL | 18/02/2016 | Kilcummin | -22.295 147.517 | 280 | 527 | - | 289 | 214 | - | - |
| 15 | AYTF16KIL | 18/02/2016 | Kilcummin | -22.295 147.517 | 280 | 408 | - | - | - | 203 | 180 |
| 16 | AYTM17DAL | 13/10/2016 | Dalby Box | -27.350 151.176 | 343 | 543 | - | 277 | 240 | - | - |
| 17 | AYTF17DAL | 13/10/2016 | Dalby Box | -27.350 151.176 | 343 | 881 | - | - | - | 360 | 503 |

<sup>a</sup>No. of unique hybrids included both the test hybrids and a range of commercial hybrids.

The trials analysed in this study formed part of the advanced yield testing (AYT) component of the pre-breeding program and contained hybrids that had already been subjected to some degree of selection for performance in preliminary trials. The 17 trials comprised 11 different sowing dates extending from 23 September to 18 February. Typical practice within the breeding program is for two trials to be grown at each site; one AYTM trial consisted of F_1_ hybrids derived from crosses between a large number of male parent lines and three female testers (i.e. B010054, B963676, and B986604), while the other AYTF trial consisted of F_1_ hybrids derived from crosses between a large number of female parent lines and two male testers (i.e. R986087-2-4-1 and R993396). The five testers varied in levels of stay-green (measured by delayed leaf senescence), plant height, and lodging (Table 2). Although the aim within each trial was to have the same number of hybrids between test lines and the respective testers, the numbers varied because of seed availability. Similarly, it was intended that each AYTM or AYTF trial series contained the same hybrids at each site in a particular year but this was also not achieved due to seed constraints. Since the trials were part of an active pre-breeding program, hybrids were added and dropped from the trial series between seasons. As a result, only a proportion of hybrids were in common between each year within the greatest coincidence being 195 in common between consecutive years.

**Table 2.** Overall Best Linear Unbiased Predictors (BLUPs) for lodging score (%), leaf senescence rating (1-9), plant height (cm), and grain yield (t ha^-1^), and the between trait correlations of the female and male testers across multiple trials.

| Tester | Tester type | Lodging score (%) | Leaf senescence rating | Plant height (cm) | Yield ($\text{t ha}^{-1}$) | Correlation coefficient between lodging and leaf senescence rating | Correlation coefficient between lodging and plant height |
| --- | --- | --- | --- | --- | --- | --- | --- |
| B010054 | Female | 9.9 | 6.4 | 120.1 | 5.13 | 0.57 | 0.42 |
| B963676 | Female | 7.7 | 6.0 | 121.3 | 5.56 | 0.34 | 0.37 |
| B986604 | Female | 7.0 | 6.2 | 116.2 | 5.53 | 0.22 | 0.32 |
| R986087-2-4-1 | Male | 10.1 | 6.4 | 125.2 | 5.22 | 0.57 | 0.25 |
| R993396 | Male | 10.8 | 6.7 | 122.0 | 5.07 | 0.58 | 0.28 |

The total number of hybrids grown per trial varied from 408 to 925 depending on season and location (Table 1), with entries of the trials including both the test hybrids detailed above and a range of commercial hybrids. All trials were arranged in a partially replicated row column trial design; on average between 20-30% of hybrids were replicated at least twice per trial, while the remaining hybrids were only grown once. All plots consisted of two rows of 5-metre length. Trials were managed according to local management practices and all were planted using a solid row configuration (i.e. no use of skip row systems). Row spacing was 0.76 or 1 metre depending on the dominant agronomic practice of the region. The trial identifiers used were a combination of trial type (“AYTM” or “AYTF”), year (such as “15” for 2015), and location (e.g. “CAP” for “Capella”).

### Phenotypes

Lodging was visually scored as the percentage of plants lodged in a plot. Leaf senescence, as a measure of stay-green, was visually rated from 1 to 9, with 1 indicating less than 10% senescent leaves and 9 indicating over 90% senescent leaves in a plot. As stay-green (i.e. delayed leaf senescence) was only expressed in trials suffering from water stress after flowering (Jordan et al. 2012), leaf senescence ratings were recorded in only 9 of the 17 trials. Plant height was measured in centimetres from the base of the plant to the tip of the panicle after maturity and was measured in 11 of the 17 trials. Only 12 of the 17 trials were harvested; the other 5 trials could not be harvested due to severe lodging just before harvest. Grains were harvested for individual plots with a small plot harvester with integrated load cells; grain yield of each plot was later converted to tonnes per hectare.

### Statistical analysis

Phenotypes were analysed using linear mixed models implemented in the asreml-R package (Butler et al. 2009) using the R software version 3.3.2 (R Core Team 2016). Genotype was included as a random effect and Best Linear Unbiased Predictors (BLUPs) for lodging, leaf senescence and plant height were predicted for all test hybrids in individual trials after accounting for spatial variation and randomisation processes used in the design. The concurrence of hybrids between locations and seasons allowed the trials to be included in multi-environment trials (MET) analysis. As there were no genotypes in common between AYTM and AYTF, MET analyses were conducted separately for the two types of trials. Factor analytic (FA) models were implemented in the MET analyses. The FA models accounted for spatial variation in individual trials and used between-trial correlations as environmental loadings through a variance matrix across trials (Smith et al. 2001). An overall BLUP for each genotype across trials was subsequently predicted.

Broad-sense heritability for each trait in individual trials was estimated according to the formula developed by Cullis et al. (2006) to account for spatial variability.

### Genotyping

DNA from mixed leaf blade tissue of two-week-old seedlings of each line was extracted using procedures described by Diversity Arrays Technology (http://www.diversityarrays.com). Samples were genotyped with the DArTseq™ platform, which represents a combination of DArT complexity reduction methods based on methyl filtration and next-generation sequencing platforms (Kilian et al. 2012). A total of 24,135 SNPs were identified for all female and male lines across three years, of which 11,435 SNPs were polymorphic. Missing data were imputed with Beagle version 4.1 (Browning and Browning 2016), with an accuracy of 94%, determined through multiple rounds of randomly masking genotypes, imputing genotype calls and correlating with genotype calls before masking. Following imputation and removal of SNPs with a minor allele count in less than 5 homozygous samples, the final marker data sets consisted of between 3,661 and 7,577 SNPs for the individual AYTM trials and between 3,290 and 3,622 SNPs for the individual ATYF trails.

### Genome-wide association study (GWAS)

GWAS for lodging was conducted separately on parent lines within each tester/trial combination, in order to avoid the confounding effect of the differences in the genetic background of the testers and the differences in environments, using FarmCPU (Liu et al. 2016). As no population structure was observed for hybrids in combination with each tester, population structure was not included as a covariate in GWAS. Parameter method.bin was set to “optimum” with the default settings of bin.size and bin.selection to optimise the possible QTN (quantitative trait nucleotide) window size and the number of possible QTNs selected into the FarmCPU model. Thirty-nine separate GWAS were conducted for the thirty-four tester/trial combinations and hybrids in combination with each of the five testers across trials (five tester populations). To integrate the GWAS output across the 39 separate analyses, a comparable method to the mpQTL approach described previously (Mace et al. 2013) was employed to calculate an integrated probability statistic, to identify regions of the genome that have a significant contribution to the variation of lodging in the tester/trial combinations studied. Specifically, the most significant marker for each tester/trial combination was selected from a sliding window of length 2 cM around each marker, based on the genetic distances calculated using the sorghum genetic linkage consensus map (Mace et al. 2009). This process was performed on each tester/trial combination separately to generate a series of probability values centred on each marker along each chromosome. The sum of these probability values across all trial combinations for each tester was then calculated, not including values where p>0.05 and plotted across the genome for each tester (Supplemental Fig. S1), with a threshold of sum(-log10(p)) >= 5.2 used to identify significance. This threshold is equivalent to a minimum of 4 trials with significance for lodging at p < 0.05 for each tester, and is more stringent than the criteria of using the Bonferroni corrected p value (0.05/Number of SNPs) for each single tester/trial combination. The peak position above the defined significance threshold for each genomic region across each tester was identified and the results combined across the five testers. Peaks within 2 cM of each other were considered as the same QTL across testers, with the peak position considered as the position with the highest sum(-log10(p)) within a 2 cM window. The confidence interval (CI) of a QTL was set to its peak position ± 1 cM. All QTL identified across the five tester populations were projected onto the sorghum consensus map (Mace et al. 2009) and visualised with MapChart v2.32 (Voorrips 2002).

QTL for lodging identified in the current study were identified as “QLDGX”, followed by chromosome number, a full-stop, and a number suffix indicating the instance of QTL occurrence on each chromosome, i.e. “QLDGX1.3” refers to the third QTL for lodging detected on chromosome 1.

### QTL haplotype identification

SNPs within the QTL confidence interval of each QTL identified were used for haplotype identification within all male lines crossed to all three female testers and all female lines crossed to the two male testers separately. There was only one or no SNP within the 2 cM confidence interval of 5 QTL for female lines and of 2 QTL for male lines; hence, these regions were extended to 4 cM to include a minimum of two SNPs. The final number of SNPs within individual QTL ranged from 2 to 93 depending on the QTL and tester population.

Given the co-ancestry of the breeding populations, the haplotypes represent regions that are identical-by-descent. Haplotypes were identified for both male and female lines using SNPs within a QTL using k-means clustering, with the number of clusters estimated by optimum average Silhouette width implemented in the “pamk” function of the “fpc” R package (Hennig 2018). Haplotype information based on all male or female lines were then extracted for each tester population.

Mean values of overall BLUPs for lodging score, leaf senescence, and plant height of each haplotype of each QTL × tester population were subsequently calculated. Analysis of variance (ANOVA) of the difference among the means of each trait for the haplotypes within each QTL/tester population was performed to test the effects of lodging QTL on leaf senescence and plant height. Haplotypes whose sample sizes were less than the number of haplotypes plus one were removed to reduce false positives. In order to determine the directions of the effects of lodging QTL on leaf senescence and plant height, mean values of leaf senescence and plant height of the two extreme haplotypes, which had the maximum and minimum average lodging scores, were calculated for each QTL/tester population. Differences in the mean leaf senescence and plant height of the two extreme haplotypes were compared to the corresponding difference in the mean lodging score for each QTL/tester population.

## Results

### Phenotypic evaluation

Mean lodging in the 17 trials varied from 2.5% to 37.3%, with heritability varying from 33.3% to 89.4% with a mean of 66.0% (Table 3). Genetic correlations of lodging between trials varied from −0.24 to 0.79, with a mean of 0.29 for AYTM and 0.39 for AYTF trials (Supplemental Table S1).

**Table 3.** Trait characteristics of hybrids grown in the 17 advanced yield testing (AYT) trials for males (AYTM) and females (AYTF) lines.

| Trial code | Trial | Sowing date | Lodging (%) |  |  | Leaf senescence |  |  | Height (cm) |  |  | Yield (t ha<sup>-1</sup>) |  |  |
| --- | --- | --- | --- | --- | --- | --- | --- | --- | --- | --- | --- | --- | --- | --- |
|  |  |  | Range | Mean | H<sup>2</sup>(%) | Range | Mean | H<sup>2</sup>(%) | Range | Mean | H<sup>2</sup>(%) | Range | Mean | H<sup>2</sup>(%) |
| 4 | AYTM15CAP | 11/01/2015 | 0 - 75 | 2.5 | 84.4 | 5 - 9 | 7.4 | 67.1 | 84 - 182 | 121.8 | 88.7 | 1.08 - 4.57 | 3.01 | 34.9 |
| 3 | AYTM15JIM | 17/12/2014 | 0 - 80 | 5.4 | 89.4 | 3 - 9 | 6.3 | 75.3 | - | - | - | 1.47 - 7.03 | 4.33 | 69.7 |
| 11 | AYTF16ORI | 15/02/2016 | 2 - 50 | 9.0 | 36.5 | - | - | - | 76 - 146 | 115.7 | 84.9 | 0.84 - 5.80 | 2.86 | 36.1 |
| 12 | AYTM16EME | 16/02/2016 | 0 - 95 | 10.7 | 60.1 | 1 - 8 | 4.2 | 52.0 | 80 - 164 | 109.3 | 85.1 | 0.00 - 8.07 | 2.63 | 54.2 |
| 8 | AYTM16WAR | 21/12/2015 | 10 - 70 | 12.6 | 33.3 | 3 - 9 | 6.0 | 74.1 | 92 - 200 | 123.5 | 90.3 | 2.64 - 11.33 | 7.25 | 63.6 |
| 10 | AYTM16ORI | 15/02/2016 | 2 - 90 | 14.7 | 62.5 | - | - | - | 76 - 160 | 109.0 | 90.2 | 0.07 - 6.39 | 2.90 | 56.9 |
| 5 | AYTM15EME | 12/01/2015 | 0 - 80 | 15.6 | 85.4 | 4 - 9 | 7.4 | 72.3 | 96 - 168 | 121.0 | 88.8 | 1.00 - 6.11 | 3.13 | 68.0 |
| 15 | AYTF16KIL | 18/02/2016 | 0 - 95 | 17.4 | 57.0 | - | - | - | 92 - 148 | 124.9 | 91.4 | 0.00 - 6.22 | 2.59 | 61.1 |
| 6 | AYTM16CRO | 23/09/2015 | 10 - 80 | 19.0 | 53.8 | - | - | - | - | - | - | 3.18 - 7.76 | 5.78 | 37.4 |
| 2 | AYTM15WAR | 3/12/2014 | 10 - 90 | 23.4 | 79.3 | - | - | - | 98 - 210 | 132.4 | 86.2 | 3.05 - 10.22 | 7.00 | 77.1 |
| 7 | AYTF16CRO | 23/09/2015 | 10 - 80 | 25.3 | 67.3 | - | - | - | - | - | - | 1.50 - 7.84 | 5.40 | 44.5 |
| 9 | AYTF16WAR | 21/12/2015 | 10 - 90 | 25.9 | 57.9 | 5 - 9 | 7.8 | 75.4 | 100 - 164 | 129.7 | 87.7 | 3.13 - 10.64 | 6.66 | 74.6 |
| 16 | AYTM17DAL | 13/10/2016 | 10 - 90 | 28.2 | 68.4 | 4 - 9 | 6.3 | 65.5 | - | - | - | - | - | - |
| 14 | AYTM16KIL | 18/02/2016 | 0 - 100 | 31.3 | 64.0 | - | - | - | 88 - 172 | 117.3 | 94.2 | - | - | - |
| 13 | AYTF16EME | 16/02/2016 | 0 - 100 | 32.2 | 67.2 | 2 - 8 | 6.0 | 45.3 | 84 - 144 | 121.6 | 87.8 | - | - | - |
| 17 | AYTF17DAL | 13/10/2016 | 10 - 90 | 33.1 | 72.9 | 3 - 9 | 6.0 | 62.5 | - | - | - | - | - | - |
| 1 | AYTM15CUR | 28/10/2014 | 10 - 90 | 37.3 | 82.7 | - | - | - | - | - | - | - | - | - |

Visual ratings of leaf senescence were recorded in 9 of the 17 trials and varied in both mean and severity of leaf senescence. Across the nine trials, the range of the leaf senescence ratings varied from 1 to 9, with mean ratings varying from 4.2 to 7.8; difference between the ratings of the most and least senescent genotypes within individual trials varied from 4 to 7 (Table 3). Heritability of the trait was generally high, varying from 45.3% to 75.4% with a mean of 65.5%. Genetic correlations of leaf senescence between trials ranged from 0.30 to 0.92, with a mean of 0.53 for AYTM and 0.39 for AYTF trials (Supplemental Table S1).

Plant height was measured in 11 of the 17 trials, with the range varying from 76 to 210 cm, and with mean plant height varying from 109 to 132.4 cm. Differences between the tallest and shortest plants within individual trials varied from 56 to 112 cm (Table 3). Heritability of plant height was very high, varying from 84.9% to 94.2%, with a mean of 88.7%. Genetic correlations of plant height between trials were also very high, varying from 0.76 to 0.97, with a mean of 0.88 for AYTM and 0.86 for AYTF trials (Supplemental Table S1).

Twelve of the 17 trials were harvested and they varied in both the range and mean of grain yield. Across the 12 trials, the range of grain yield varied from 0 to 11.33 t ha^-1^, with mean yield varying from 2.59 to 7.25 t ha^-1^ (Table 3). Heritability of grain yield varied from 34.9% to 77.1%, with a mean of 56.5%. Genetic correlations of yield between trials varied from −0.07 to 0.71, with a mean of 0.20 for AYTM and 0.11 for AYTF trials (Supplemental Table S1).

Weak to moderate correlations between overall BLUPs for lodging and leaf senescence, and between overall BLUPs for lodging and plant height, were observed, ranging from 0.22 to 0.58, ranged from 0.25 to 0.42 (Fig. 1). In individual trials, correlations between lodging and leaf senescence, between lodging and plant height were also weakly to moderately positive for all five tester populations (Supplemental Fig. S2).

**Fig. 1.**
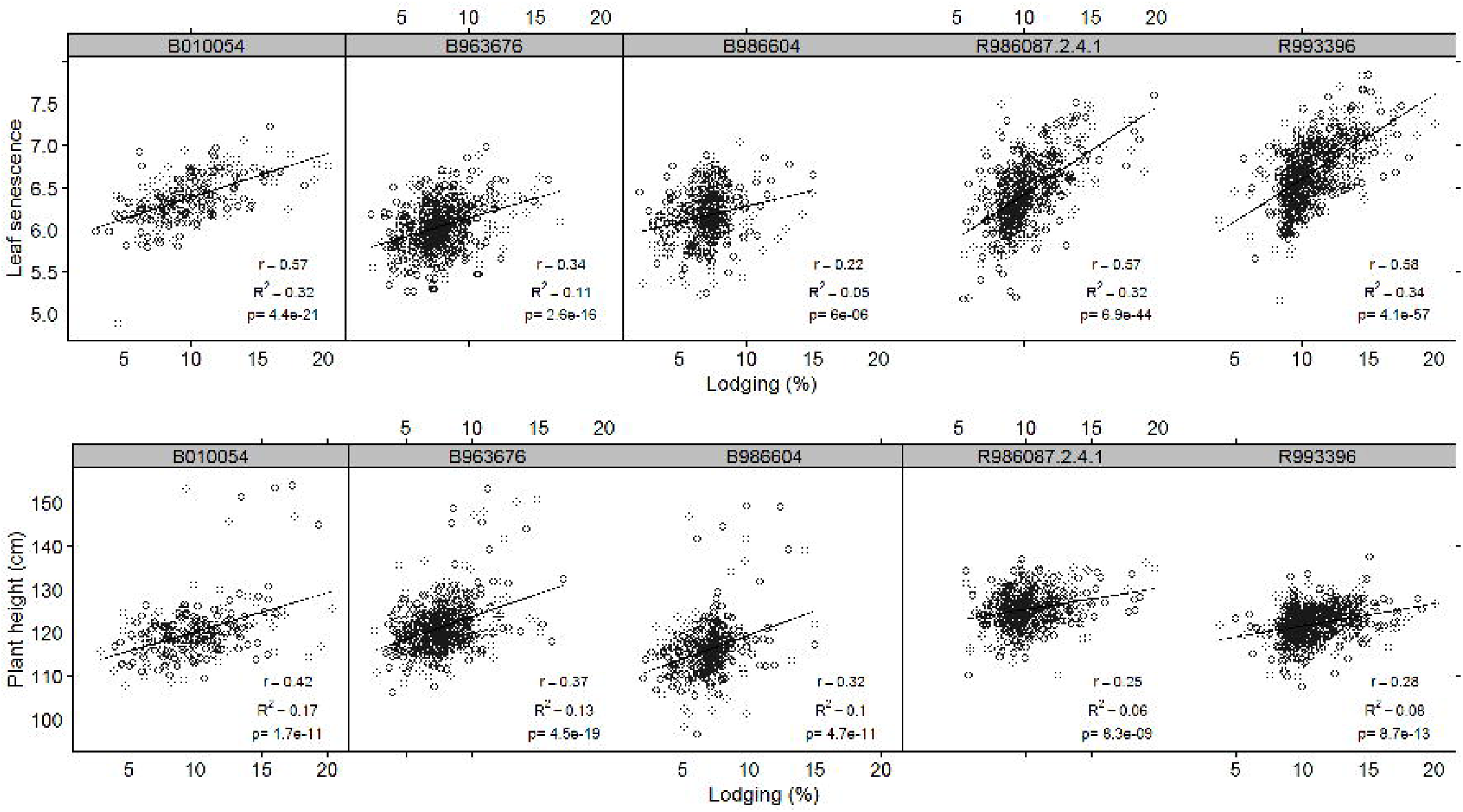
Correlations between overall BLUPs for leaf senescence, plant height, and lodging by tester. B010054, B963676, B986604, R986087-2-4-1, and R993396 are the five tester parents of the hybrids grown in the 17 yield testing trials.

Between-trait correlations of hybrids varied between the testers (Table 2; Fig. 1; Supplemental Fig. S2). Comparisons could only be made within female or male testers but could not be made between male and female testers because even though they were grown at the same locations, the male and female tester hybrids were grown in different trials and there were differences in the genetic variance of the traits in the different parental pools. However, the same general trends were observed. Testers with low breeding values for leaf senescence in both female and male trials showed low correlations between leaf senescence and lodging (Table 2; Fig. 1; Supplemental Fig. S2). The female tester with the lowest breeding value for height showed the lowest correlation between hybrid height and lodging. In contrast, within the male testers the line with the higher breeding value for height (R986087-2-4-1) displayed a lower association between height and lodging.

### QTL identification

Across the 39 tester/trial combinations, a total of 213 unique QTL for lodging were detected. These QTL were distributed across all 10 chromosomes. Each individual QTL was detected in multiple tester/trial combinations, with 75% of the QTL identified in at least five tester/trial combinations (Supplemental Table S2).

The number of QTL identified per tester population differed largely, with the least number of QTL (i.e. 59 and 64 respectively) detected for the R986087-2-4-1 and R993396 hybrids of the AYTF trials and the most number of QTL identified for the B963676 hybrids in the AYTM trials (176 QTL). The majority of the lodging QTL were in common across tester populations, with 61% (i.e. 129) displaying significant effects in at least two tester populations (Supplemental Table S2). Of the lodging QTL which were identified in only one population, the majority (i.e. 60/84) were identified in the B963676 hybrid trials.

### Association of leaf senescence and plant height with lodging QTL

The QTL that affected lodging showed consistent directions of effect for both plant height and leaf senescence. In both the male and female populations, the QTL haplotypes with high lodging tended to have high leaf senescence. Similarly, the QTL haplotypes with high lodging tended to be taller. The 213 QTL by 5 testers produced 1065 instances where the difference between the extreme lodging haplotypes could be compared for the other two traits. For leaf senescence in 808 of 1065 instances, the high lodging haplotype corresponded with the high senescence haplotype. For plant height in 799 of 1065 instances, the high lodging haplotype corresponded with the tall haplotype. The direction and significance of these haplotype differences are shown in Supplemental Table S2.

Only 16 lodging QTL were not significantly associated with either plant height or leaf senescence in at least one tester. In addition, 6 lodging QTL were associated with leaf senescence or plant height but showed the opposite haplotype effects (i.e. high lodging haplotypes had lower leaf senescence or were shorter). These 22 QTL were defined as non-pleiotropic effect QTL (npQTL) (Fig. 2; Supplemental Table S2).

**Fig. 2.**
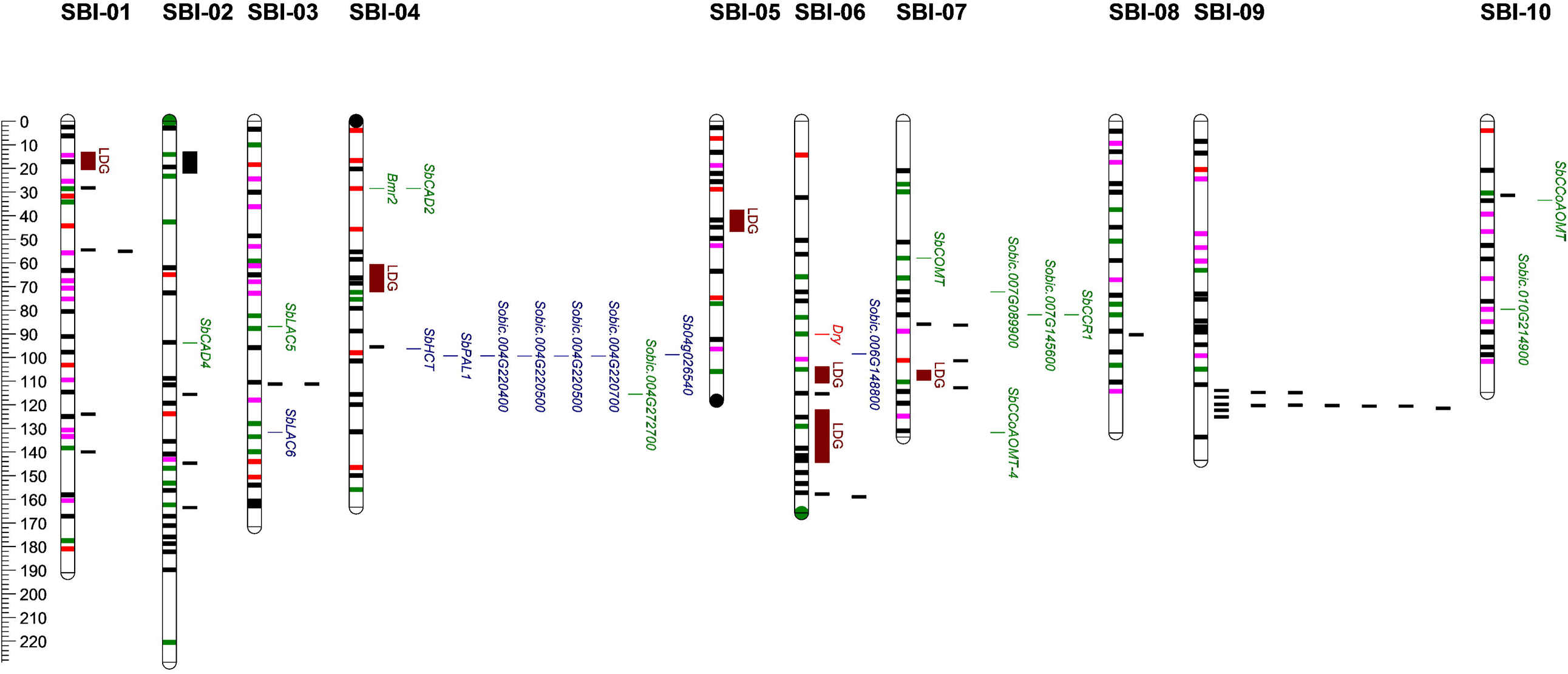
Projection of the 213 lodging QTL identified across testers onto the sorghum consensus genetic map and their co-localisation with previously identified QTL for lodging and stalk rots and genes in lignin biosynthesis pathway. The 213 lodging QTL identified as horizontal segments on each chromosome and colour-coded as follows: lodging QTL that significantly affect leaf senescence (green), lodging QTL that significantly affect plant height (pink), lodging QTL that significantly affect both leaf senescence and plant height (black), QTL did not affect either leaf senescence or plant height (red). Brown vertical bars to the right of each chromosome marked as “LDG” represent the locations of QTL previously detected for lodging (Kebede *et al.*, 2001; Murray *et al.*, 2008; Srinivasa Reddy *et al.*, 2008); the black bars indicate the location of a QTL previously reported for charcoal rot and/or Fusarium stalk rot (Srinivasa Reddy et al. 2008; Adeyanju et al. 2015). The location of genes in the lignin biosynthesis pathways that co-located with lodging QTL detected in this study are highlighted in green to the right of each chromosome, and those that did not co-locate with lodging QTL are highlighted in blue (Bout and Vermerris 2003; Saballos et al. 2009, 2012, Sattler et al. 2009, 2017, Walker et al. 2013, 2016, Jun et al. 2017, 2018; Wang et al. 2017). The location of the recently cloned *Dry* gene is highlighted in red (Zhang et al. 2018).

### Co-localisation with previously identified QTL for lodging and stalk rots

Six of the seven previously identified QTL contributing to lodging resistance (Kebede et al. 2001; Murray et al. 2008; Srinivasa Reddy et al. 2008) were able to be projected onto the sorghum genetic linkage consensus map and all 6 overlapped with lodging QTL identified in this study (Fig. 2). These six QTL were identified in mapping populations of between 93 and 176 individuals and hence had large confidence intervals.

Two previous studies detected 34 unique QTL for resistance to charcoal rot and/or Fusarium stalk rot (Srinivasa Reddy et al. 2008; Adeyanju et al. 2015), of which 12 overlapped with lodging QTL detected in this study (Fig 2). Eleven of these 12 overlapping lodging QTL were associated with leaf senescence and/or plant height, with only one of the charcoal rot QTL co-locating with the npQTL for lodging.

### Co-localisation with genes for stem composition

A limited number of genes for stem composition have been identified and functionally validated in sorghum. Based on phylogenetic relationships with functionally confirmed genes in sorghum and other species, predicted translational protein functions, enzymatic activity assay, and gene expression patterns, 21 candidate genes in the lignin biosynthesis pathways have been identified in sorghum (Bout and Vermerris 2003; Saballos et al. 2009, 2012, Sattler et al. 2009, 2017, Walker et al. 2013, 2016, Jun et al. 2017, 2018; Wang et al. 2017). Additionally, the major morphological *D* gene, which changes the structure of the sorghum pith, was recently cloned (Zhang et al. 2018). Each of these genes was compared to the location of the QTL for lodging identified in this study.

Twelve of the 21 genes in the lignin biosynthesis pathway in sorghum were located within the lodging QTL identified in this study, representing a significant enrichment (χ^2^ test, p value = 0.008) (Fig. 2). To confirm this enrichment, comparison was made to multiple sets of 21 randomly selected sorghum house-keeping genes identified from *Arabidopsis thaliana* (Scheideler et al. 2002) and these were found not to be significantly enriched within the lodging QTL identified in this study (χ^2^ test, p value = 0.07, average of 20 independent random samplings).

The recently cloned *Dry* gene controls the stem juice content and the composition and structure of the stem pith, and influences the water content of the stem (Zhang et al. 2018). For this reason, the gene is likely to affect stem strength. It was found to be co-located with one of the QTL associated with leaf senescence, QLDGX6.9.

## Discussion

Sorghum crops experiencing drought stress during grain filling often lodge, resulting in substantial yield losses. Lodging can be affected by many factors such as plant stature, stay-green and stalk rots. Plant height may further influence lodging by changing the mechanical stress experienced by the stem. If stalk rotting pathogens are present and environmental circumstances are favourable, lodging is further exacerbated by the degradation of the stem. Direct selection for resistance to stalk rots has not proven to be effective in reducing lodging susceptibility. To date, selection for lodging resistance has been based on selection against lodging susceptibility and to some extent selection for stay-green (Jordan et al. 2012). A major challenge to this approach is that genotypes that are more susceptible to lodging also tend to have higher yield potential and hence are more likely to suffer carbohydrate shortages when carbon supply is restricted by drought. As a result, selection for lodging resistance constrains genetic gain for grain yield and vice versa. Little is known about the genetic architecture of the trait due to its complexity. Here, with a view to improving the efficiency of selection for lodging resistance, we present the largest investigation of the genetic architecture of lodging in sorghum to date. The results of this study have revealed the complexity of the genetic architecture of the trait with over 200 QTL identified. Of these QTL, over 90% were also associated with differences in plant height and leaf senescence.

### Lodging is under complex genetic control

In this study, we conducted GWAS across 39 separate tester/trial combinations involving 2308 unique elite F_1_ hybrids in 17 pre-breeding trials across 9 locations in 3 growing seasons. The populations explored variation in both the male and female parental pools. We identified 213 QTL for lodging, which was equivalent to an average of 5.6 QTL per tester/trial combination. Three quarters of the QTL were detected in at least five and 91 QTL in more than 10 tester/trial combinations (Supplemental Table S2). The large number of QTL identified in this study demonstrates that lodging is a highly quantitative trait, and is one of the most complex traits reported to date.

All previous QTL mapping studies for lodging resistance in sorghum were conducted in small bi-parental populations (93-176 individuals) in a limited number of environments (1-3). These studies have identified a total of 7 QTL, on average 2.3 QTL/study, none of which overlapped between studies (Kebede et al. 2001; Murray et al. 2008; Srinivasa Reddy et al. 2008). However, 6 of the 7 QTL projected onto the sorghum consensus map (Mace et al. 2009) overlapped with QTL identified in the current study (Fig. 2). It was very likely that large population sizes, diverse genetic background of breeding lines, and the large number of growth environments contributed to the identification of more QTL for lodging in this study.

### Carbohydrate remobilisation and plant height are the major drivers of lodging

Leaf senescence rated at maturity in sorghum subjected to water stress during grain filling provides a measure of the degree of remobilisation required to fill the developing grain (Borrell and Hammer 2000). The degree of senescence is related to the imbalance between the demand and supply of carbohydrate within the plant. Remobilisation of carbohydrate reserves from the stem and leaves ultimately results in tissue death and predisposes the plant to lodging. The degree of lodging can be further modified by constitutive factors of the stem and by pathogen invasion (Henzell et al. 1984). In this study, we found that the lodging score was positively correlated (p<0.001) with leaf senescence, across environments and genetic background of testers (Table 2; Fig. 1; Supplemental Fig. S2A). Sixty-nine percent of the 213 lodging QTL identified showed a significant positive association between lodging score and leaf senescence (Supplemental Table S2). This clearly suggests an important role of carbohydrate remobilisation in lodging. This is in line with the earlier studies (Rosenow 1980; Henzell et al. 1984; Rosenow and Clark 1995) that showed increased lodging was associated with increased leaf senescence.

This and other studies led to the adoption of the stay-green trait (delayed leaf senescence) as an additional protective trait in Australia and the USA, where it has led to significant increases in grain yield as well as reduced lodging (Henzell et al. 1992; Henzell and Hare 1996; Rosenow et al. 1997; Jordan et al. 2012). The results from both our study and previous studies indicate that the stay-green trait, a delayed leaf senescence, is a very useful trait in reducing stalk lodging.

This study also detected a significant positive association between lodging score and plant height regardless of the environments and genetic background of testers (Fig. 1; Supplemental Fig. S2B). A positive association between lodging and plant height has also been reported by earlier studies (Esechie et al. 1977; Henzell et al. 1984; Srinivasa Reddy et al. 2008; Gomez et al. 2017, 2018). In agreement with the phenotypic association between plant height and lodging score, the haplotype ANOVA results revealed that 66% of the 213 lodging QTL identified in this study significantly affected lodging susceptibility with taller plants being more susceptible to lodging (Supplemental Table S2). However, similar to stay-green, differences in the association between plant height and lodging were observed amongst the testers (Table 2; Fig. 1). Testers with more stay-green were less susceptible to negative effects of increased plant height on lodging. In fact, the male tester with a lower leaf senescence value (R986087-2-4-1) produced hybrids with higher average height but showed a lower association between height and lodging. Similar protective effects of stay-green were observed when the female testers were compared. Compared to B010054, the less leaf senescent female tester B963676 that also produced taller hybrids on average showed a lower association between height and lodging.

We also investigated the impact of yield potential on lodging because high yielding genotypes tend to remobilise more stem reserves for grain filling, which potentially causes more lodging. Our data suggest that yield potential is an important factor in lodging. Overall BLUPs for lodging were predicted to get a measure of lodging propensity for each hybrid and single trial BLUPs for grain yield were predicted for each hybrid in individual trials. We calculated the correlation between overall BLUPs for lodging and single trial BLUPs for yield for trials where lodging did not occur and observed significant, positive correlations in 24 of the 45 tester/trial combinations and significant, negative correlations in only 4 combinations (Supplemental Table S3).

### The impact of stalk rotting pathogens

Stalk rots can also accelerate lodging of infected plants. Very few studies have identified QTL for resistance to stalk rot in sorghum to date. Twelve of the 34 unique stalk rot resistance QTL previously detected (Srinivasa Reddy et al. 2008; Adeyanju et al. 2015) overlapped with lodging QTL in this study, 11 of which were associated with leaf senescence and/or plant height (Fig. 2). Given that *Fusarium* spp. and *Macrophomina phaseolina* are necrotrophs (Dodd 1980) and the observation of the substantial association with remobilisation and/or plant height, it seems likely that classical resistance genes are of limited importance in the selection for enhanced lodging resistance.

### The impact of stem composition

Twelve of the 21 genes involved in lignin biosynthesis (Bout and Vermerris 2003; Saballos et al. 2009, 2012, Sattler et al. 2009, 2017, Walker et al. 2013, 2016, Jun et al. 2017, 2018; Wang et al. 2017) were located within genomic regions associated with lodging in this study (Fig. 2), representing a significant enrichment (χ^2^ test, p value = 0.008). Lodging resistant sorghum generally have more lignification, both in the number of cells with lignified walls and in the degree of lignification in the epidermis, sub-epidermis, and vascular bundles (Schertz and Rosenow 1977). Similar results have been previously observed in maize, with reduced lignin content in the stem resulting in increased stalk lodging (Sun et al. 2018). More recently, Zhang et al. (2020) reported that downregulation of a maize gene, *stiff1*, resulted in thicker stalk cell walls with increased cellulose and lignin, which improved stalk strength and reduced lodging.

Previous studies have reported that sorghum lines with both increased (e.g. *SbMyb60* overexpression plants) and reduced lignin content (e.g. *bmr* lines) can provide the same level of or higher resistance to *Fusarium* spp. and *Macrophomina phaseolina* than sorghum lines with normal lignin content (Funnell-Harris et al. 2010, 2014, 2017, 2018, 2019). This might be due to the inhibition the infection of stalk rotting pathogens by cell-wall-bound phenolic compounds from monolignol biosynthesis that can induce defensible pathways (Funnell-Harris et al. 2017). Therefore, the role of lignin in lodging resistance may act in two ways, either by providing mechanical support to the stem (Bashford et al. 1976) or by restricting the infection of stalk rotting pathogens.

The recently cloned *Dry* gene controls the stem juice content and the composition and structure of the stem pith, and influences the water content of the stem (Zhang et al. 2018). We found that the *Dry* gene co-located with one of the QTL associated with leaf senescence (QLDGX6.9). Using markers within the *Dry* gene, four haplotypes were identified and they differed in lodging severity (data not presented). For the above reasons, we believe that the *Dry* gene is likely to affect stem strength and thus lodging, which may or may not be associated with stalk rots as reported by Funnell-Harris et al. (2016) that both susceptible and resistant sweet sorghum lines to *F.* spp. and *M. phaseolina* had been identified (Funnell-Harris et al. 2016).

These two co-localisations with lodging QTL suggest stem composition plays an important role in lodging, and could be a useful target for selection.

### Implications for breeding and future research

The identification of 213 QTL in this study demonstrates that lodging is a highly quantitative trait with multiple QTL each of small effect.

Lodging is affected by many traits with confounding effects. However, one primary cause of lodging is the imbalance between the supply and demand of carbohydrate during grain filling. A breeding program selecting for lodging resistance *per se* is likely to be of limited effectiveness because of the trade-off with yield. Our study showed that there is a strong association between grain yield potential and lodging susceptibility (Supplemental Table S3). This suggests that sorghum breeders must be cautious when selecting directly for lodging resistance as it is likely to significantly constrain genetic gain for yield. The results support the current practice of direct selection for increased yield and indirect selection for lodging resistance by selecting for delayed leaf senescence (stay-green). In addition to its role in reducing lodging, the integrating stay-green phenotype has been shown to be associated with increased yield in most Australian sorghum environments (Henzell et al. 1992; Henzell and Hare 1996; Borrell et al. 2000, 2014b; Jordan et al. 2012) and the USA (Rosenow et al. 1997).

Our study confirmed the undocumented observation by Australian sorghum industry personnel that increased plant height is associated with increased propensity to lodging (Table 2; Fig. 1; Supplemental Fig. S2B). However, as with yield, sorghum breeders need to be cautious when selecting for lodging resistance through the selection for short plants because taller plants tend to have higher potential grain yield (Jordan et al. 2003). Our research indicates that the traits associated with the integrating stay-green phenotype provides protective effects, weakening the association between plant height and lodging (Table 2; Fig. 1). Therefore, there would appear to be opportunities to increase genetic yield gain by increasing height while reducing lodging susceptibility through selection for traits associated with non-senescence.

Although stalk rots can promote the development and the severity of lodging, previous studies suggest stalk rot susceptibility is affected by source-sink relationships and considered to be a problem under post-anthesis stresses (Dodd 1980; Henzell et al. 1984; Rosenow 1984; Rosenow and Clark 1995). Consistent with these early studies, we found that 12 of the 34 QTL for stalk rot resistance co-located with lodging QTL in this study, 11 of which were related to leaf senescence and/or plant height (Fig. 2). Direct selection for resistance to stalk rots has not been proven to be effective in reducing lodging susceptibility. Our data indicates that classical resistance genes to stalk rots are likely to be of limited importance in breeding for increased lodging resistance.

Co-localisations of the genes for stem composition with lodging QTL in the study highlight the importance of stem strength and composition in lodging resistance. More research is required to investigate the relationship between morphological, mechanical, and composite traits of the stem and lodging resistance. The identification of traits related to physical stem strength that can improve resistance to lodging may prove to be a more effective and efficient way by reducing the dependency of screening lodging in drought conditions during grain filling.

## Conclusions

The detection of 213 QTL clearly demonstrates that water stress induced lodging is a complex trait in grain sorghum. The majority of these QTL were associated with leaf senescence and/or plant height, which suggests that lodging in grain sorghum in Australia is mainly driven by the source-sink relationship of carbohydrate and plant height. Sorghum breeders need to be cautious when selecting for both lodging resistance *per se* and short plants because of the likely negative consequences on grain yield. Instead, our study supports selecting for stay-green (delayed leaf senescence) to reduce lodging susceptibility. Additionally, our data indicated that the stay-green phenotype displayed a protective effect, weakening the positive association between lodging susceptibility and plant height. Therefore, the combination of selection for both delayed leaf senescence and increased plant height could allow for both enhancing genetic yield improvement and reducing susceptibility to lodging. Additionally whilst our study indicated that resistance to lodging is unlikely to benefit from the selection for classical resistant genes to stalk rots, selection for stem composition could play an important role in enhancing lodging resistance.

## Declarations

### Availability of data and material

The data and material used in this study are availability upon request to the corresponding author.

### Code availability

Not Applicable.

### Author contributions

X.W., D.J., and G.H. conceived the project. X.W., D.J., E.M., A.C., and Y. T. contributed to the development of the analytical framework and the synthesis of the results. X.W. and C.H. conducted the statistical analyses. X.W. conducted GWAS analysis. X.W. wrote the manuscript. All authors read and revised the manuscript.

## Acknowledgements

We would like to acknowledge the sorghum pre-breeding team at Hermitage Research Facility for support of the field trials.

## Supplemental Information

**Fig. S1 Diagram of the method to integrate the GWAS output across the 39 separate analyses, a comparable method to the mpQTL approach describe by Mace *et al.* (2013).**

To identify genomic regions that have a significant contribution to the variation of lodging, an integrated probability statistic was calculated by summing the -log10(p) of the p values (5^th^ to 10^th^ columns) of the most significant SNP for each tester (B010054 in this example) and trial combination from a sliding window of length 2 cM around each SNPs (A). The process was performed on each tester/trial combination separately to generate a series of probability values centred on each SNP along each chromosome. The sum of these probability values across all trial combinations for each tester was then calculated, excluding values where p>0.05 and plotted across the genome for each tester (B). The position of the SNP in the centred of the 2cM sliding window (shaded in grey) is highlighted in orange and the corresponding sum(-log10(p)) within the window highlighted in green; values highlighted in yellow indicates the most significant p values for the corresponding tester/trial combination within the 2cM window, whereas blue highlighted values indicates p values that are < 0.05 but not included in the calculation of sum(-log10(p)).

**Fig. S2 Correlations between BLUPs for lodging and leaf senescence (A), lodging and plant height (B) in individual trials by tester.**

Table S1 Between-trial genetic correlations of lodging, leaf senescence, plant height, and grain yield.

Table S2 Details of lodging QTL detected across tester populations.

Table S3 Correlation between overall BLUPs for lodging score and single trial BLUPs for grain yield in all 38 trials by tester. Mean lodging score, mean leaf senescence rating, and mean yield of the tester/trial combinations are also presented. Five trials, AYTM15CUR, AYTM16KIL, AYTM17DAL, AYTF16EME, and AYTF17DAL, were not harvested due to lodging; therefore, yield BLUPs in these trials could not be predicted and correlations in these trials could not be calculated.

