## Supplemental Fig. S1, Supplemental Fig. S2 for "Large scale genome-wide association study reveals that drought induced lodging in grain sorghum is associated with plant height and traits linked to carbon remobilisation"

### Slide 1
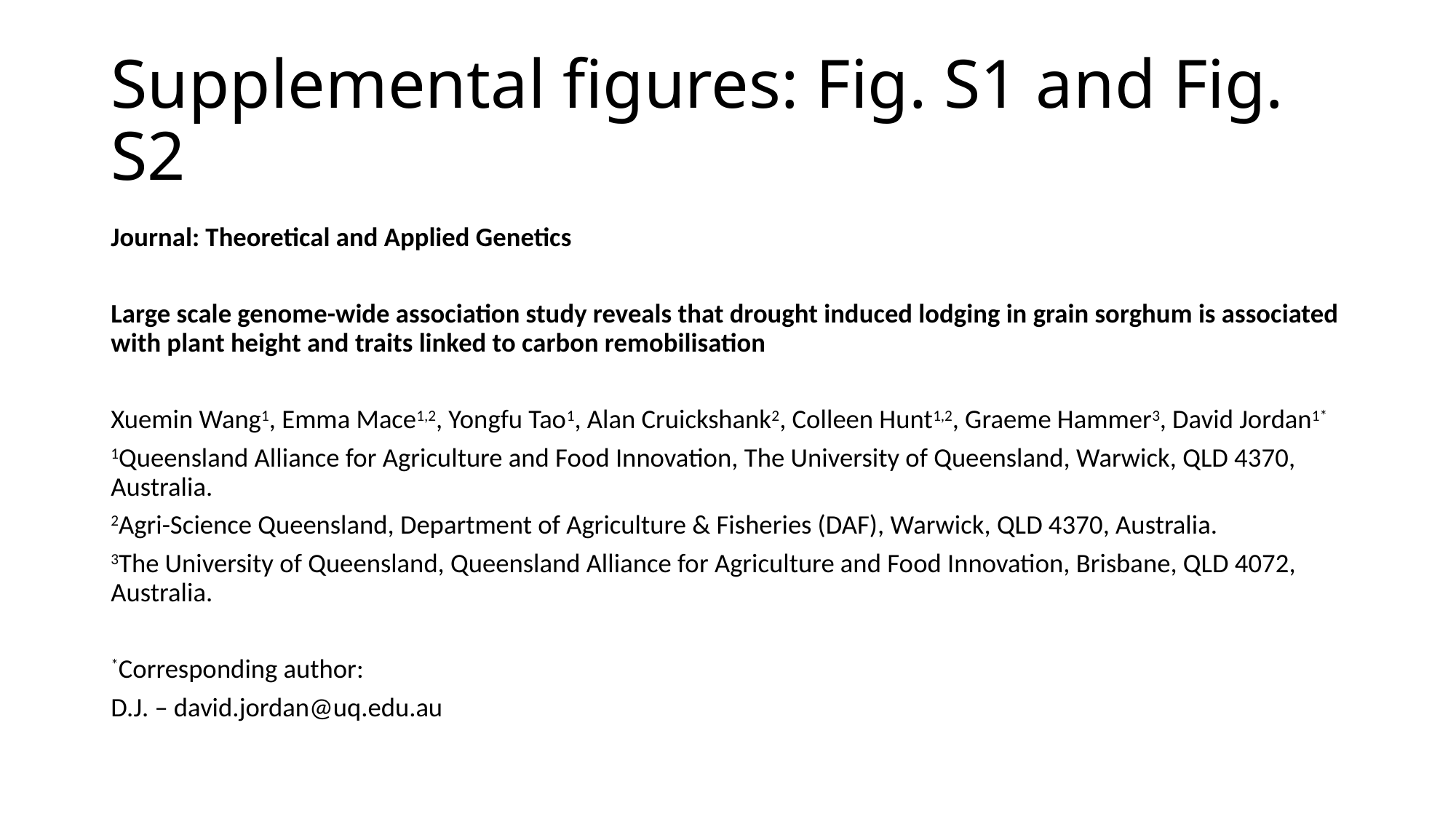

#### Supplemental figures: Fig. S1 and Fig. S2
Journal: Theoretical and Applied Genetics
Large scale genome-wide association study reveals that drought induced lodging in grain sorghum is associated with plant height and traits linked to carbon remobilisation
Xuemin Wang^1^, Emma Mace^1,2^, Yongfu Tao^1^, Alan Cruickshank^2^, Colleen Hunt^1,2^, Graeme Hammer^3^, David Jordan^1*^
^1^Queensland Alliance for Agriculture and Food Innovation, The University of Queensland, Warwick, QLD 4370, Australia.
^2^Agri-Science Queensland, Department of Agriculture & Fisheries (DAF), Warwick, QLD 4370, Australia.
^3^The University of Queensland, Queensland Alliance for Agriculture and Food Innovation, Brisbane, QLD 4072, Australia.
^*^Corresponding author:
D.J. –

### Slide 2
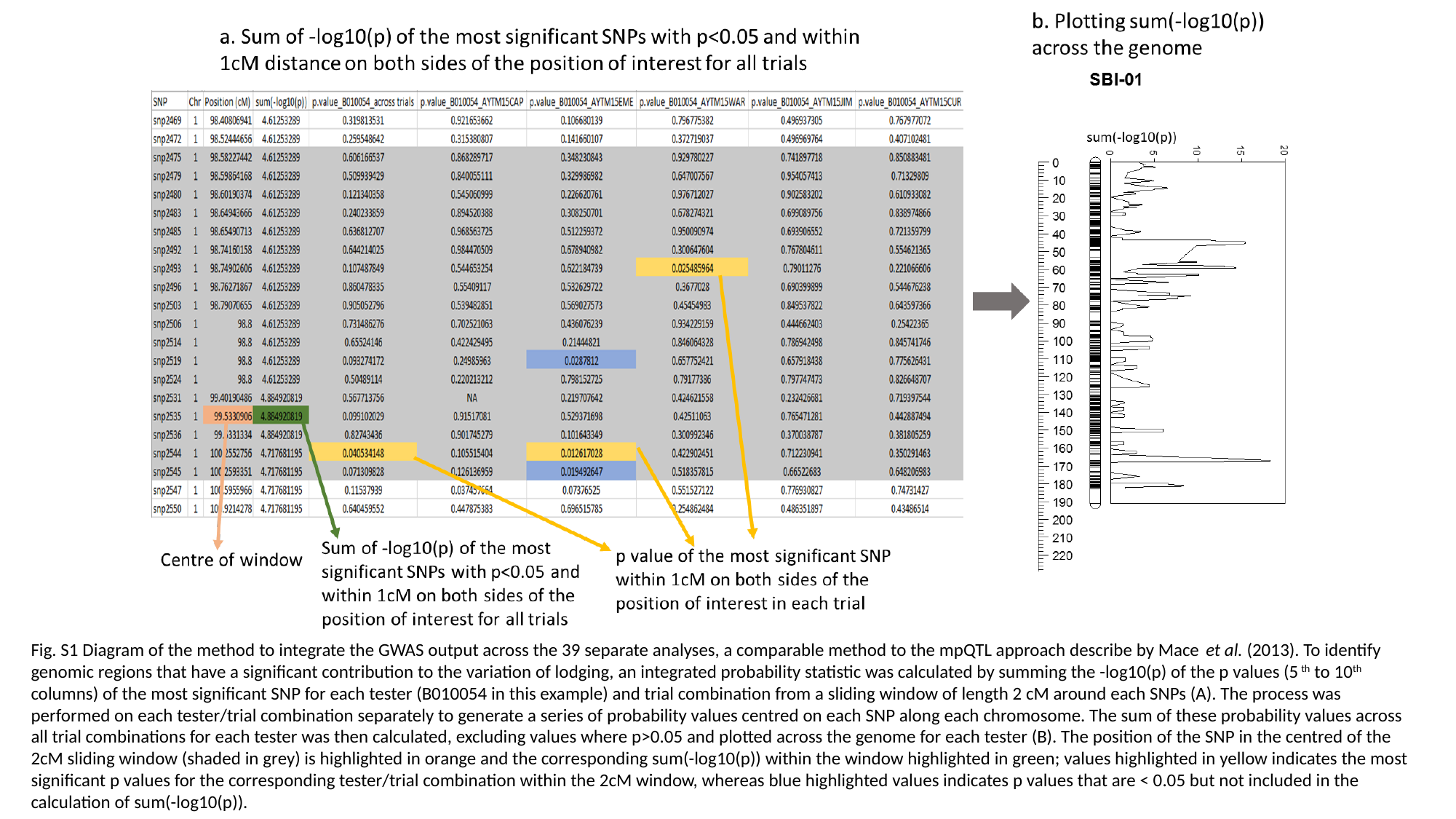

Fig. S1 Diagram of the method to integrate the GWAS output across the 39 separate analyses, a comparable method to the mpQTL approach describe by Mace et al. (2013). To identify genomic regions that have a significant contribution to the variation of lodging, an integrated probability statistic was calculated by summing the -log10(p) of the p values (5^th^ to 10^th^ columns) of the most significant SNP for each tester (B010054 in this example) and trial combination from a sliding window of length 2 cM around each SNPs (A). The process was performed on each tester/trial combination separately to generate a series of probability values centred on each SNP along each chromosome. The sum of these probability values across all trial combinations for each tester was then calculated, excluding values where p>0.05 and plotted across the genome for each tester (B). The position of the SNP in the centred of the 2cM sliding window (shaded in grey) is highlighted in orange and the corresponding sum(-log10(p)) within the window highlighted in green; values highlighted in yellow indicates the most significant p values for the corresponding tester/trial combination within the 2cM window, whereas blue highlighted values indicates p values that are < 0.05 but not included in the calculation of sum(-log10(p)).

### Slide 3
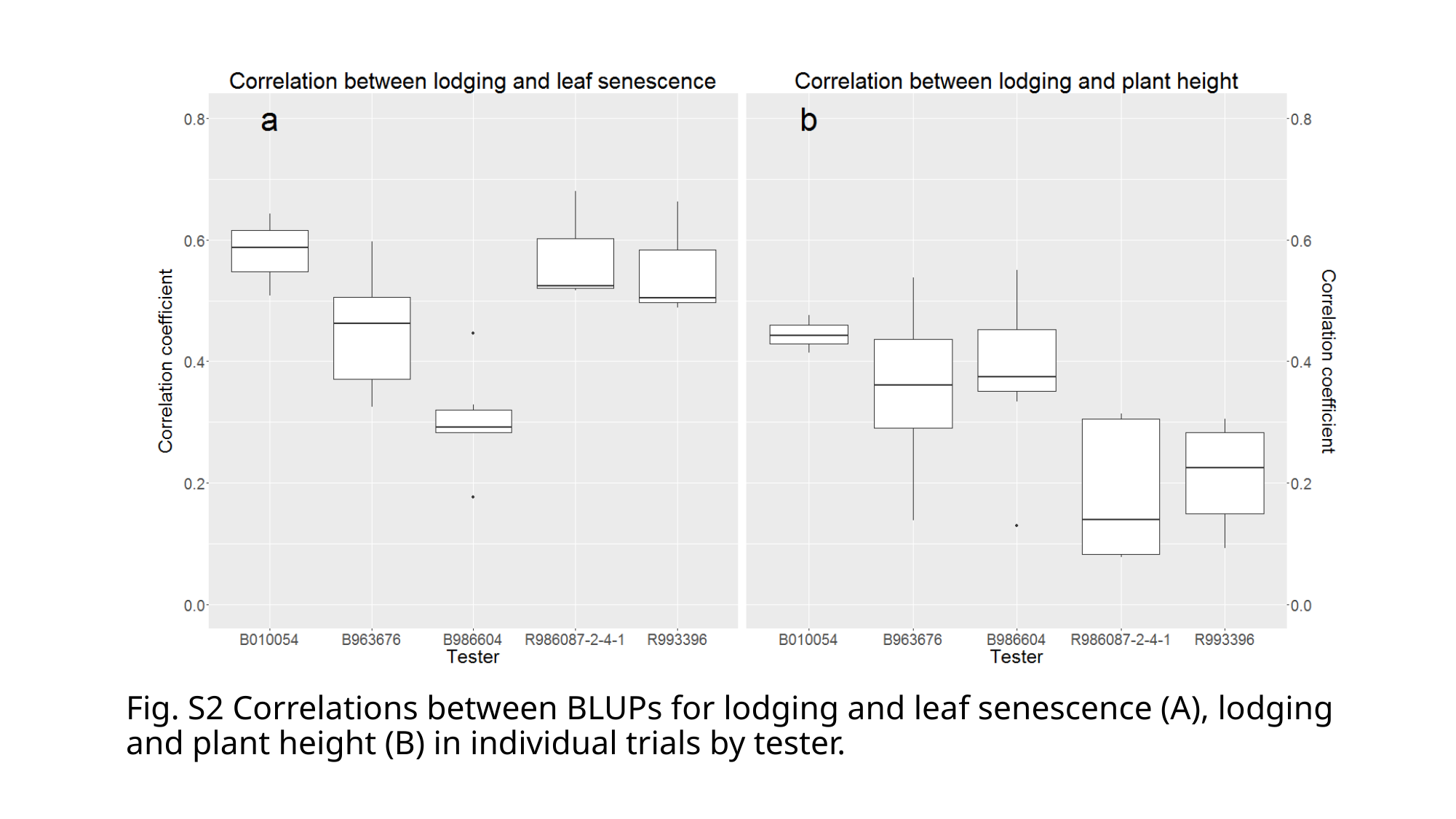

#### Fig. S2 Correlations between BLUPs for lodging and leaf senescence (A), lodging and plant height (B) in individual trials by tester.
